# Histopathological analysis of selected organs of *Oreochromis niloticus* due to sub-lethal industrial effluents exposure

**DOI:** 10.1101/2022.01.13.476240

**Authors:** Ghazala Jabeen, Sarwat Ishaq, Farkhanda Manzoor

## Abstract

On a daily basis, our environment is exposed to tons of a composite of industrial effluents, which has a negative impact on commercial fish production and, as a result, on humans. Present study was designed to evaluate the acute, sub-chronic, and chronic toxicity of a composite of raw industrial effluent from Sunder Industrial Estate in the freshwater fish *Oreochromis niloticus* biosystem by investigating at the histopathological changes in different organs such as heart, kidney, and muscle after exposure. Fish was exposed to 1/3^rd^, 1/5^th^ and 1/10^th^ of predetermined LC_50_. Significant histopathological alterations in heart (myocarditis, pericardium bending and lifting) kidney (renal tube degeneration, glomerulus structural alteration and necrotic proximal tubule) and muscle (inflammation, atrophy and tumor) were observed in treated groups. After the sub-lethal exposure histological alteration index (HAI) was highest in chronic group as compared to the acute and sub-chronic group as HAI _group D_> HAI _group C_> HAI _group B_. Moreover physicchemical parameters of water were found to be out of the range of the APHA standard approach.

## Introduction

The addition of undesired substances into water bodies causes changes in the aquatic system’s physical, chemical, and biological features, resulting in ecological imbalance (1). Industrial effluents contribute significantly to water contamination, endangering aquatic plants and animals (2). Because Industrial effluents contain variety of heavy metals (3), phenols (4) and genotoxins (5) etc. The aquatic flora and wildlife, notably fish, are steadily declining as a result of pollution (6).

As a result of untreated wastewater, fish have pathogenic abnormalities. Under chronic exposure conditions, these exposure concentrations may not even be harmful for the affected species, but they could include an impact on the internal structures and functions, affecting the performance of vital processes and functions such as environmental resistance and competitive stress, growth and reproduction (7).

Even at very low concentrations, long-term exposure to water contaminants has been found to cause histological, biochemical alterations and morphological changes in fish tissues and organs, which may have a major impact on fish quality and perhaps fish survival rate in water ((8) and (9)). In laboratory and field research, histopathological changes have been used as indicators in monitoring the fish health exposed to toxicants. Changes in these organs are frequently simpler to detect than functional changes, and they serve as early warning signs of animal illness (10).

Histopathological study has proved to be an extremely sensitive metric for detecting cellular changes in target organs (11). Histopathological biomarkers have a variety of advantages when it comes to detecting long-term damage in aquatic species, and they’re probably the easiest chronic biomarkers to apply in aquatic biology. Biomarkers of contamination exposure in species of fish are crucial indicators if we are to maintain a profitable fishery and a healthy product for men consumption and health (12).

Fish is a significant source of animal protein, particularly for people on a limited budget (13). Pollution and changes in their surroundings make fish particularly vulnerable. In polluted aquatic ecosystem, fish health is used as an indicator ((14), (15)). Before causing changes in fish behavior or appearance, pollution’s harmful effects may manifest at the cellular or tissue level (16). Examination of fish histology could be used as a bio-monitoring approach for water contamination, according to (17). Several investigations have documented histological alterations in fish exposed to effluents, in recent years ((18) and (19)).

Our environment is exposed to tons of industrial effluents on daily basis. According to several studies, a mixture of industrial effluents causes considerable changes in fish ((20), (21) and (22)). As a result, the current study attempted to investigate the variation produced by a composite of industrial effluents at different concentrations in *Oreochromis niloticus* vital organs (kidney, heart, and muscle), as well as the interaction between effluent toxicity and histological abnormalities.

## Materials and Methods

The study area was Sundar Industrial Estate, one of Lahore’s most rapidly developing and polluted industrial regions. The industrial region covers 1750 acres of land and is home to 400 major and medium sized businesses such as engineering, chemical, paint, pharmaceutical, textile, and food businesses (Figure #1). The study region is located between 31.2883° N and 74.1739° E. Water from these industries is discharged into the land on a regular basis. The people who live in the vicinity of these locations are at risk of being polluted by the environment.

**Figure 1:**
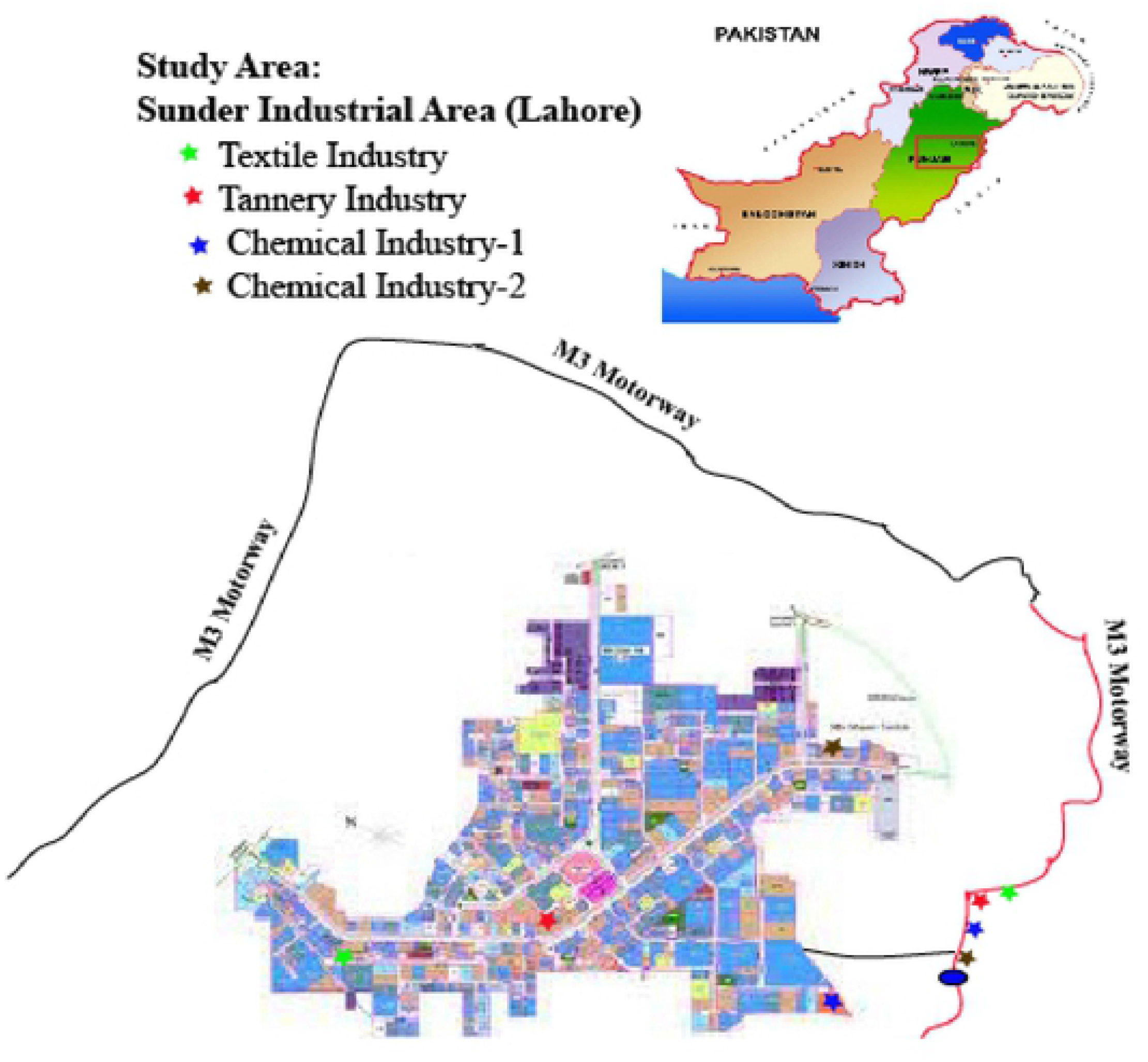
Geographical map of study area

As stock solutions, raw unfiltered effluents from Sunder Industrial Estate were collected from the points of discharge of the Textile, Tannery, and Chemical industries (1.5 L). All of the samples were taken in triplicate. “Standard Methods for Examination of Water and Wastewater” was used to record physico-chemical parameters (23). Following that, required amounts were measured into calibrated glass containers and built up to the needed concentration using the toxicity evaluation techniques provided by (24).

The experiment was conducted at three durations of 3-day, 15-day and 180-day. According to the study design followed by the guidelines of OECD, 2019, healthy juveniles were exposed to sub-lethal doses (1/3rd, 1/5th and 1/10th concentrations) of 96-hour LC_50_ value. In 3 days, 15 days, and six month static/renewal bioassay for 1/3^rd^, 1/5^th^ and 1/10^th^ of LC_50_ exposure doses and control exposures (0.00%) devoid of effluent mixes were employed. The 1/3^rd^, 1/5^th^, and 1/10^th^ concentrations of 96-hour LC_50_ value from exposures to predefined levels (v/v) of a composite of industrial effluents were 6%, 3.6%, and 1.8%, respectively. Healthy juveniles of (n=100), *Oreochromis niloticus* with an average weight of (10±2g) and length (3±0.5 inch) were transported in oxygenated waterproof bags to the Aquaculture Facility in the Department of Zoology, Lahore College for Women University, Lahore, Pakistan, Fish were kept in tanks filled with dechlorinated tap water for seven days to allow them to adjust to their new surroundings. Fish were fed a 40% crude protein diet at 3% body weight on a daily basis, with uneaten food being siphoned off on a frequent basis to prevent the accumulation of metabolites. Following acclimatization, fish were exposed to 1/3rd, 1/5th, and 1/10th of the LC50 concentrations of 6%, 3.6% and 1.8% respectively.

Animal health and behavior were monitored by observing the sub-lethal clinical signs exhibited by the fish during the exposure experiment in order to improve the analysis to examine the industrial effluent toxicity and reduce the animal suffering by following the recommendations on the identification, assessment, analysis, and use of clinical signs as humane endpoints for experimental animals used in safety evaluation for mammalian studies (OECD, 2000; OECD, 2019; EPA, 2007, 2012). The fish were monitored for consecutively 8 hours immediately after the start of exposure (day 0-1). Afterwards, 2 observations were conducted during the first 24 hours of the experiments at an interval of 3 hours. From the 2nd day to the end of the experiment, control and treated fish were inspected twice per day (preferably early morning and late afternoon to best cover the 24-hour periods). Fish behavior, mortalities and visible abnormalities were also noted like jumping, restlessness, loss of equilibrium, clustering around aerators, inverted positions such as head up or down, loss of pigmentation, scale and fin erosion, exophthalmia and floating at surface or sinking were recorded and photographed.

At the end of experimental durations, a natural anesthetic, clove oil extracted from the distilled stems, leaves and flowers of the clove tree, *Syzgium aromaticum* was used at a dose of 50 μl/L (AVMA Guidelines, 2013) for the euthanasia of fish as an approach to follow humane endpoints. The animals were euthanized after 30 minutes of sampling from the water aquaria. The animals when reached the endpoint criteria were randomly sampled, dissected and analyzed for the histological changes in the fish organs. The kidneys, hearts, and muscles of the fish were removed carefully and preserved in 10% neutral-buffered formalin before being dehydrated in ascending degrees of alcohol and cleaned in xylene. Using a Euromex Holland microtome, the preserved tissues were soaked in paraffin wax and sliced into 4-6m thick pieces. The Harris Hematoxylin and Eosin technique was used to stain the sections (25). Then these sections were viewed under a microscope and images were captured by using a microscopic camera. At last, the sections produced from the controlled fish were compared. Figure 2 depicts the overall experiment.

**Figure 2:**
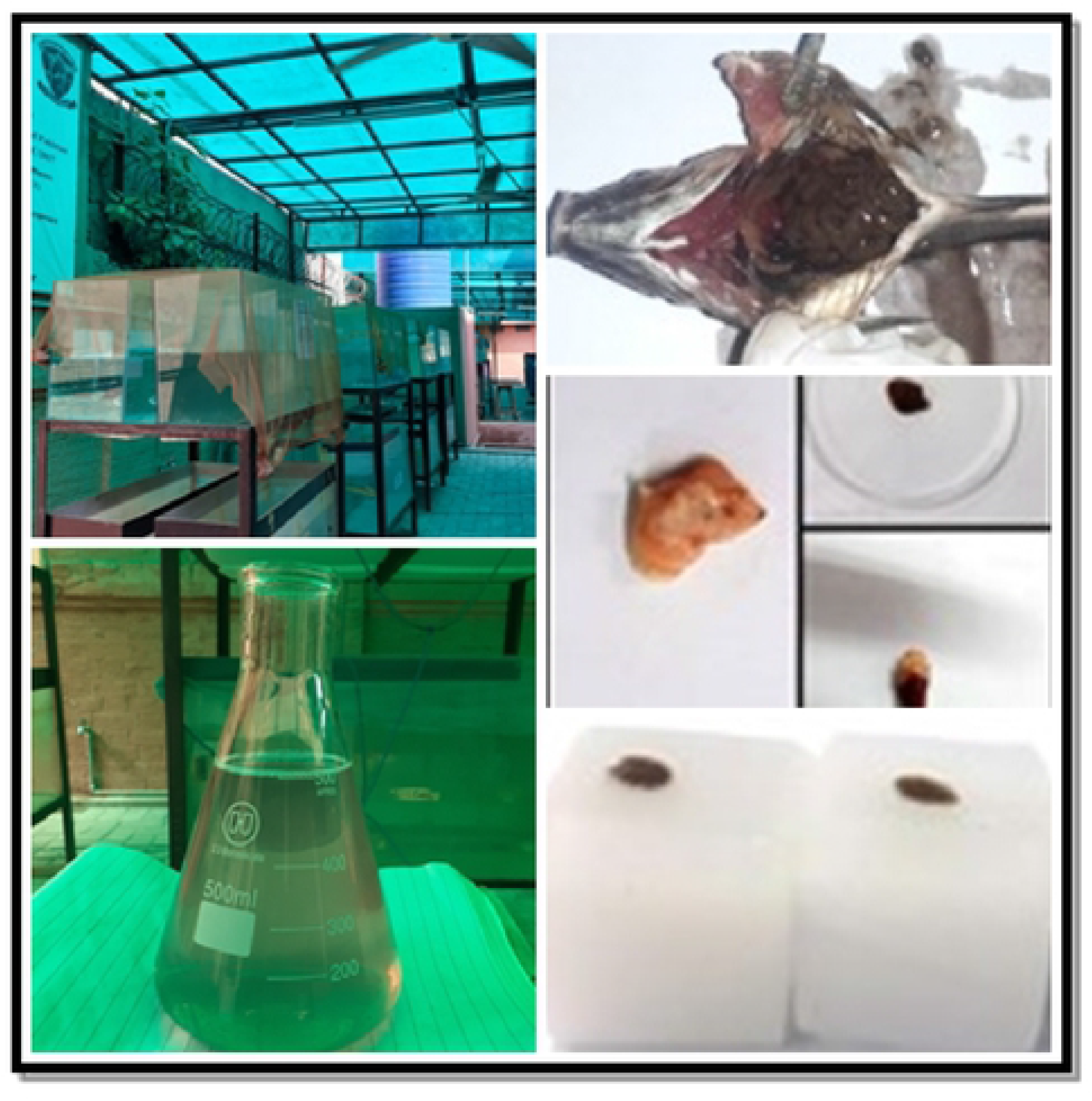
Overall representation of experiment

## Results and Discussion

Present study was conducted to evaluate the toxic impact of industrial effluents of Sunder Industrial Estate. Physico-chemical analysis revealed that these effluents were highly toxic, not only to aquatic organisms but the entire web chain including human beings who are primary consumer of fish. Aquatic species have been deemed an outstanding and cost-effective model system because of their ability to concentrate, store, and metabolize water-borne contaminants (26) for analyzing the carcinogenic, mutagenic and toxic potential of pollutants (27) and (28). Fish exposed to chemical pollutants had a variety of lesions in various organs (29). Muscle (30), kidney (11), and heart (31) are good organs to examine histologically to check how pollution affects them.

Analysis of physico-chemical parameters, temperature, pH, Free CO 2, alkalinity as CaCO3, Total alkalinity, total hardness, chloride, electrical conductivity, total dissolved solids (TDS) and salinity of control and treated groups is presented in Table 1. The mean values of all water quality variables recorded during the present investigation revealed that they all were significantly higher than the stander values. Whereas highest fluctuations were recorded in TDS content and alkalinity as CaCO_3_ of control and treated fish. TDS was recorded in leather industry as 4959.33±675.17 mgL −1 that was significantly higher than that of control group with a mean value of 356.867±303.86 mgL −1. Effluents from leather industry showed complete absence of free CO2. The mean electrical conductivity EC (μScm-1) ranged from 5835±794.21 to 1629±20.53. Highest values of EC were recorded in leather industry and lowest in chemical industry-I respectively.

**Table 1:**
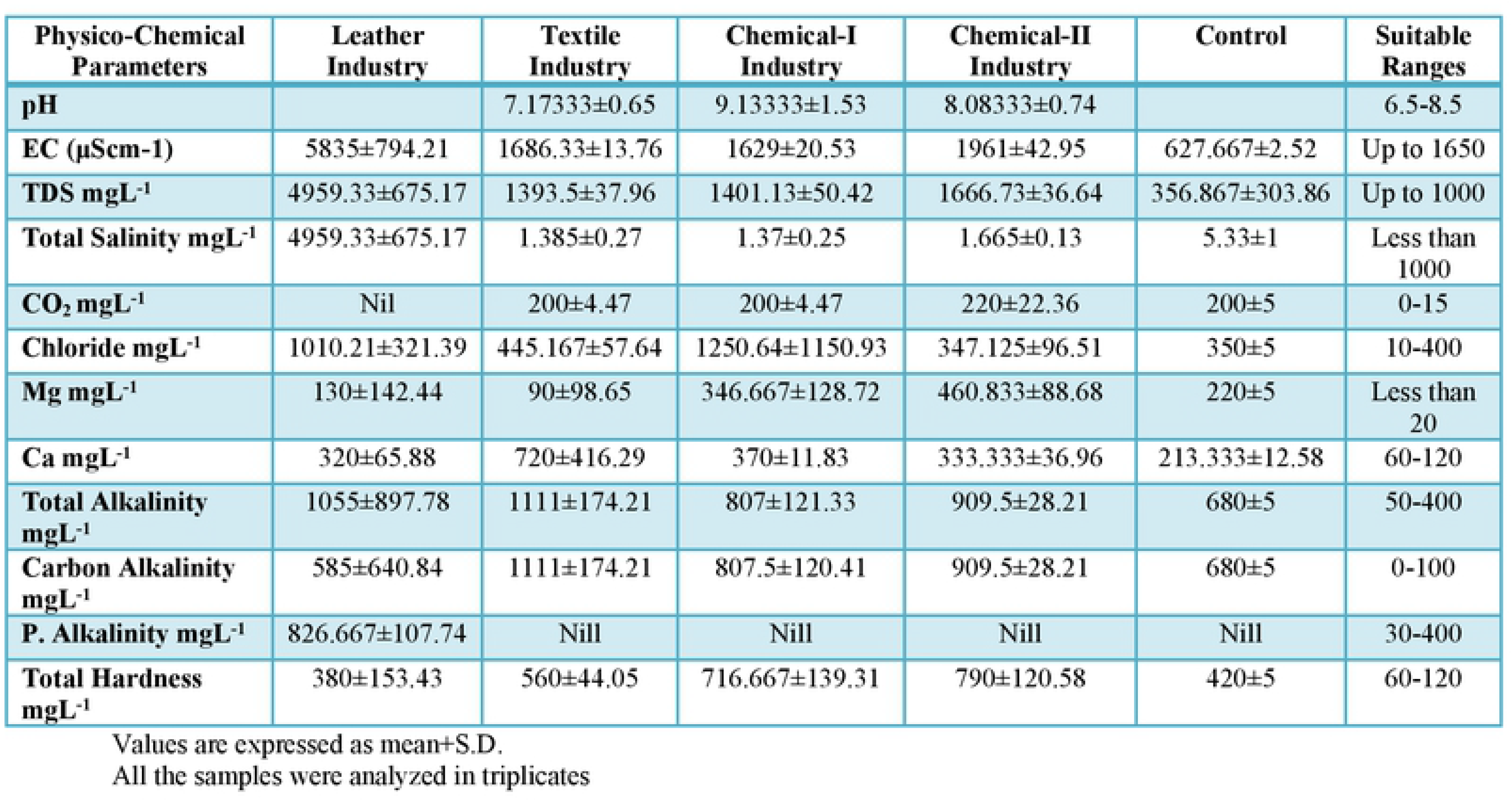
Physio-Chemical Parameter of Industrial Effluents of Sunder Industrial Estate,Lahore.

As fish were divided in two groups: control (Group D1) and treated groups (Groups D2, D3 and D4) and control group was taken as reference group. Control fish exhibited normal behavior as compared to the abnormal behavior of fish exposed to sub-lethal doses (1/3rd, 1/5th and 1/10th concentration of LC50) of industrial effluents. Fish in reference groups showed an ideal and normal physical appearance and behavior. No skin damage and scales loss was observed. Fish maintained its normal swimming position and opercular movement during the entire study period. While in treated group, a remarkable change in behavior was observed. Fish spent most of the time at the bottom with gross pathologies like exophthalmia due to outward bulging of eyes, bleeding, hemorrhages and loss of vision. Red spots were observed around the gills. Reduced swimming activity and darting was also noticed in treated group fish. Fish was lethargic after exposure to industrial effluents. Inflammation, swelling, clogging of eyes, exophthalmia and excessive mucus secretion were observed in fish exposed to sub-lethal levels (1/3rd, 1/5th and 1/10th concentration of LC_50_ of industrial effluents collected from all the sampling stations (Figure 2.1 A-F).

**Figure 2.1.**
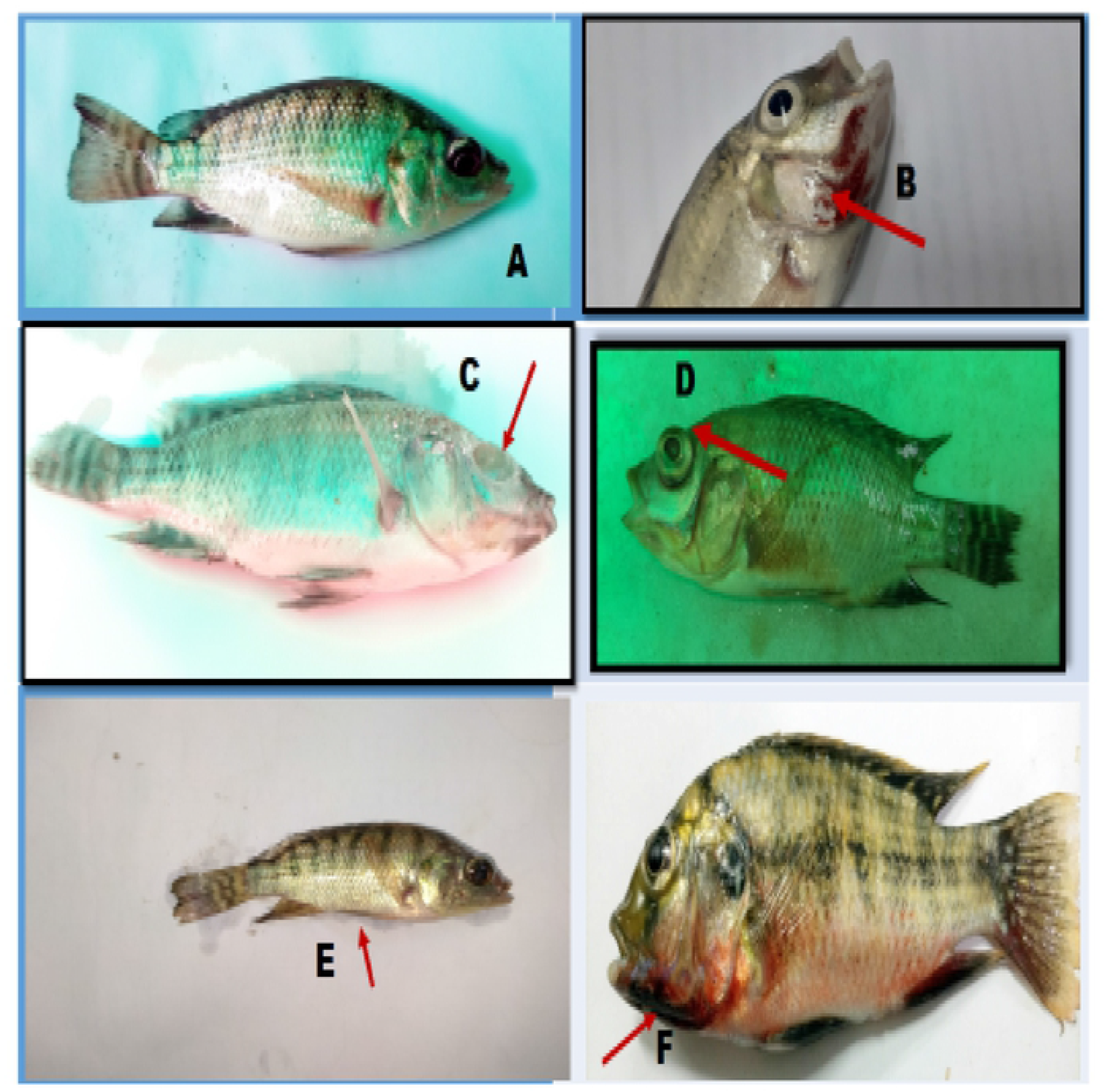
(A-F). Normal features of Oreochromis nilotocus (A); Inflammation and swelling (B); Clogging of eye and loss of vision (C); Exophthalmia (D); skeletal deformity and curvature (E) and Hemorrhage (F) due to sub-lethal industrial effluent exposure.

In test organisms, industrial effluents produce severe histological damage in the gill, kidney, liver, and gut. Furthermore, (32) revealed that malfunctions in fish general health conditions owing to direct discharge of effluents into water channels has an effect on fish general health. In the current investigation, histological changes in normal organs subjected to various concentrations of LC_50_ of industrial effluents from the Sunder Industrial Estate are depicted in Figures 3–5, indicating mild to severe abnormalities, i.e. odema, hemorrhage, inflammation, atrophy, leukocytes infiltration, glomerular variations as well as alteration in proximal tubules, hyperplasia and tumor.

**Figurer 3:**
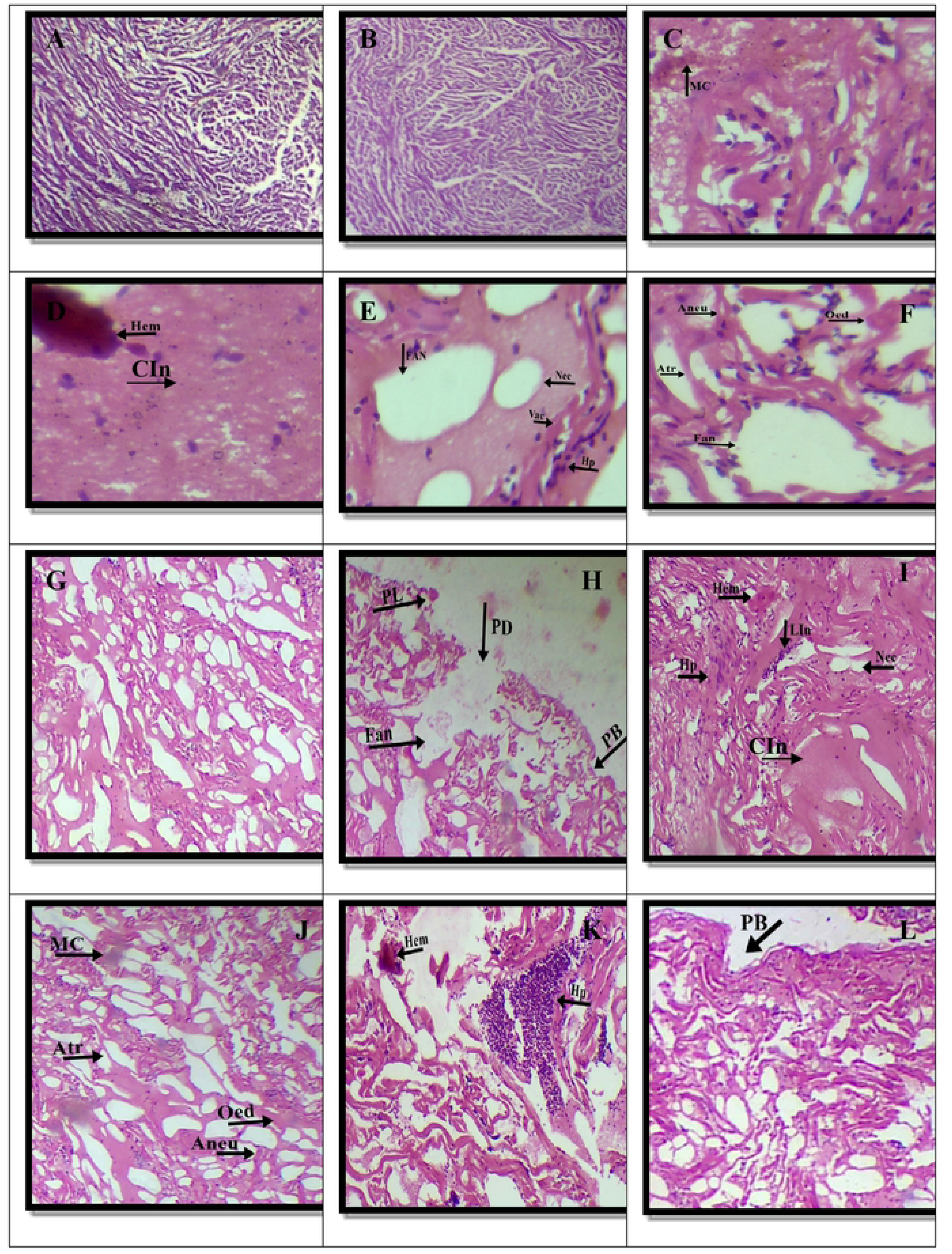
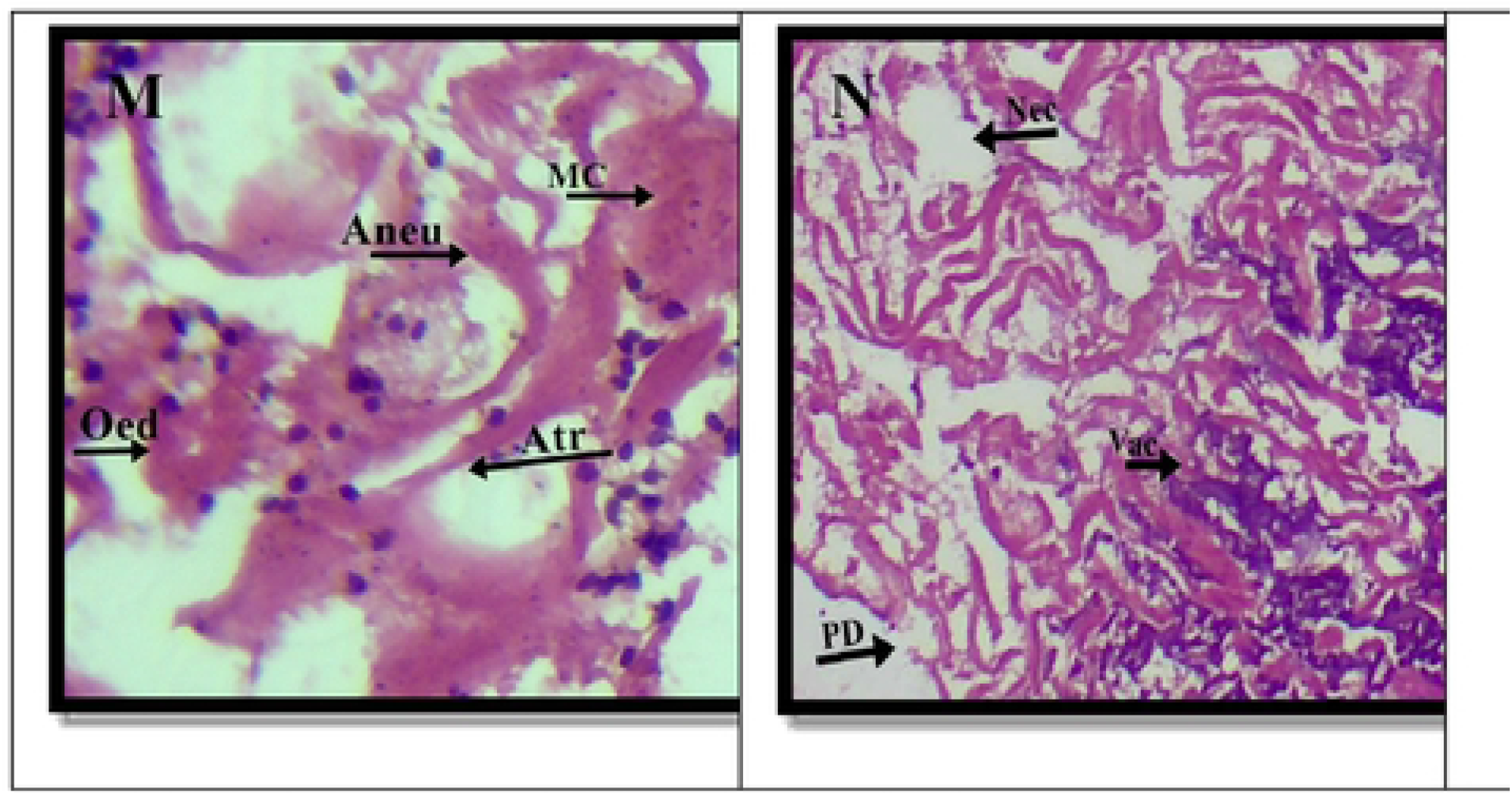
Histopathological alteration of transversely paraffin sectioned of Heart of *Oreochromis niloticus* electron micrographs (10×&40×) (H&E). (A&B) Control, (C-F) 1/3^rd^ of LC_50_, (G-J) 1/5^th^ of LC_50_, (K-N) 1/10^th^ of LC_50_. Normal architecture of muscle tissues were observed in the control fishes. **Abbriviations are as=** Hem=Hemorrhage, FAN= Focal area of necrosis, Oed=Oedema, PB=Pericardium bending, PL=Pericardium lifting, Aneu=Aneurysm, Atr=Atrophy, PD=Pericardium Damage, Vac=Vacuolar degeneration, Nec=Necrosis, HP=Hyperplasia, LI=Leucocytes Infiltration and MC=Myocarditis

**Figure 4:**
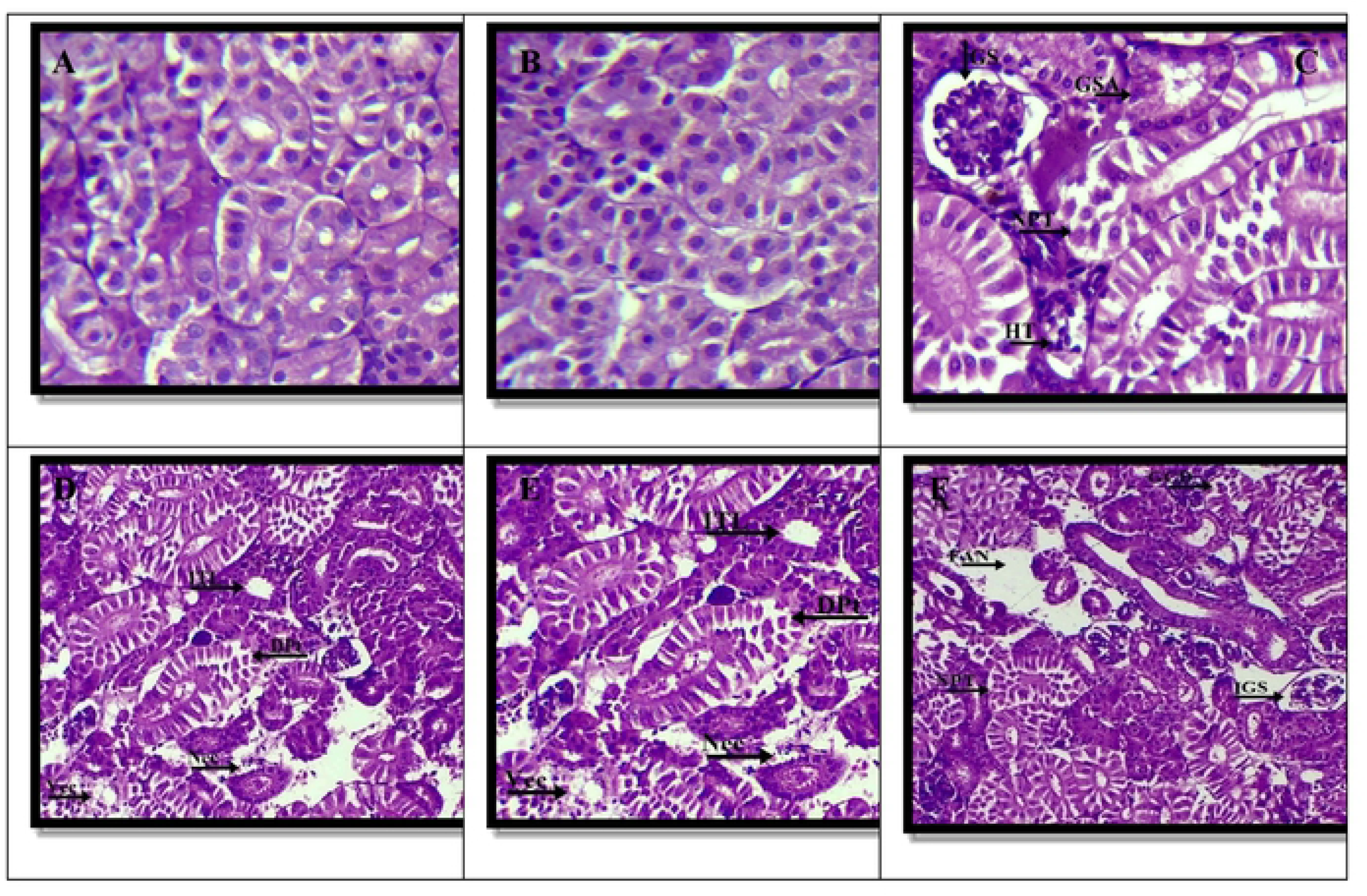
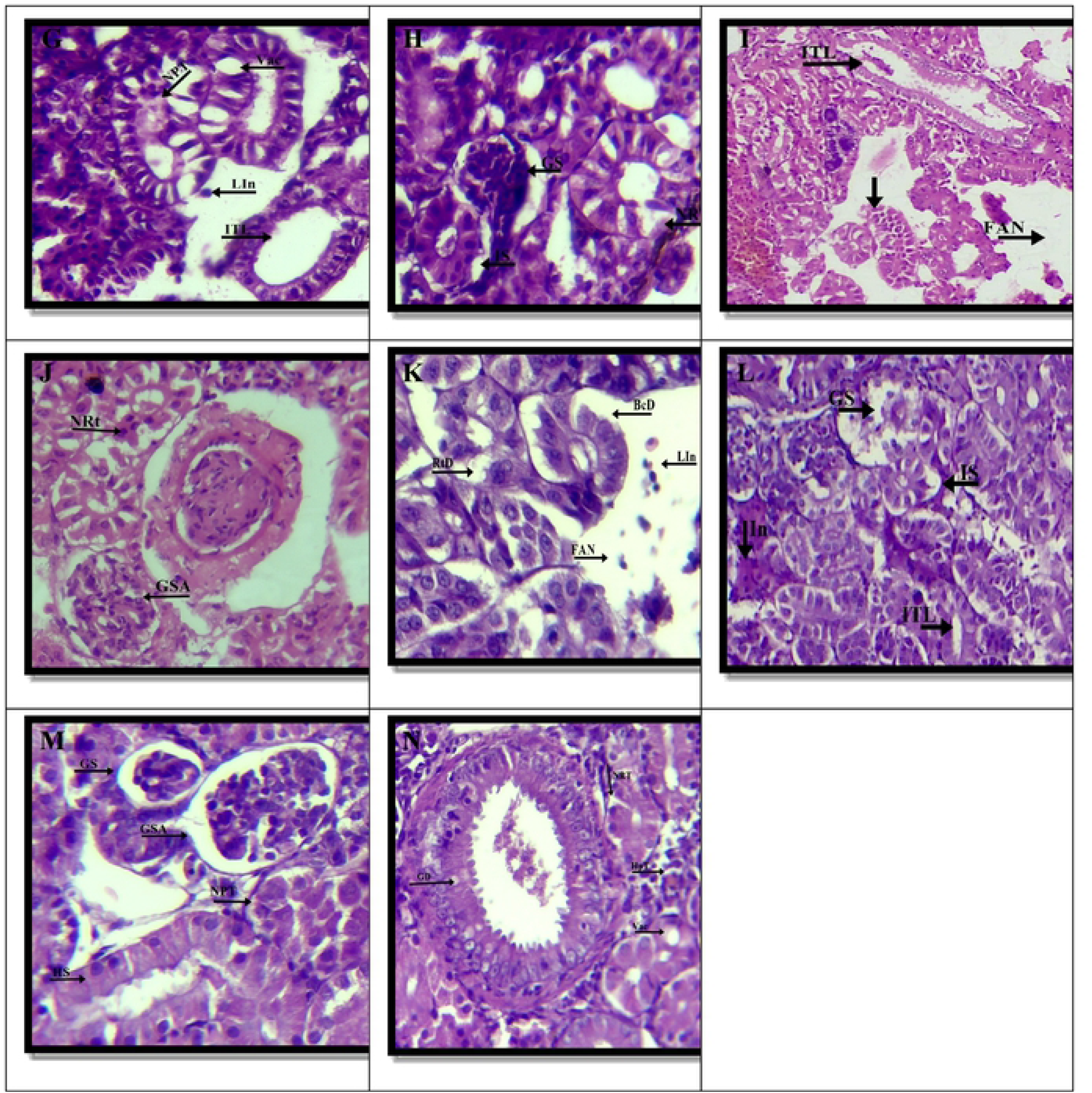
Histopathological alteration of transversely paraffin sectioned of Kidney of *Oreochromis niloticus* electron micrographs (10×&40×) (H&E). (A&B) Control, (C-F) 1/3^rd^ of LC_50_, (G-J) 1/5^th^ of LC_50_, (K-N) 1/10^th^ of LC_50_. Normal structure of kidney tissues was observed in the control fishes. **Abbriviations are as=** Hem= Hemorrhage, FAN= Focal area of necrosis, GD= Glomerulus arteries degeneration, NPt= Necrotic proximal tubule, IGS=Increase Glomerulus space, GSA=Glomerulus structural alteration, BcD= Bowman’s capsule disintegration, GS=Glomerulus shrinkage, NRT=Necrotic renal tubule, HS=Hydropic swelling, ITL=Increase tubular lumen, Vac=vacuolar degeneration, HpT=Hematopoietic tissue Necrosis, RtD=Renal tubule degeneration and Lin=Leukocyte infiltration

**Figure 5:**
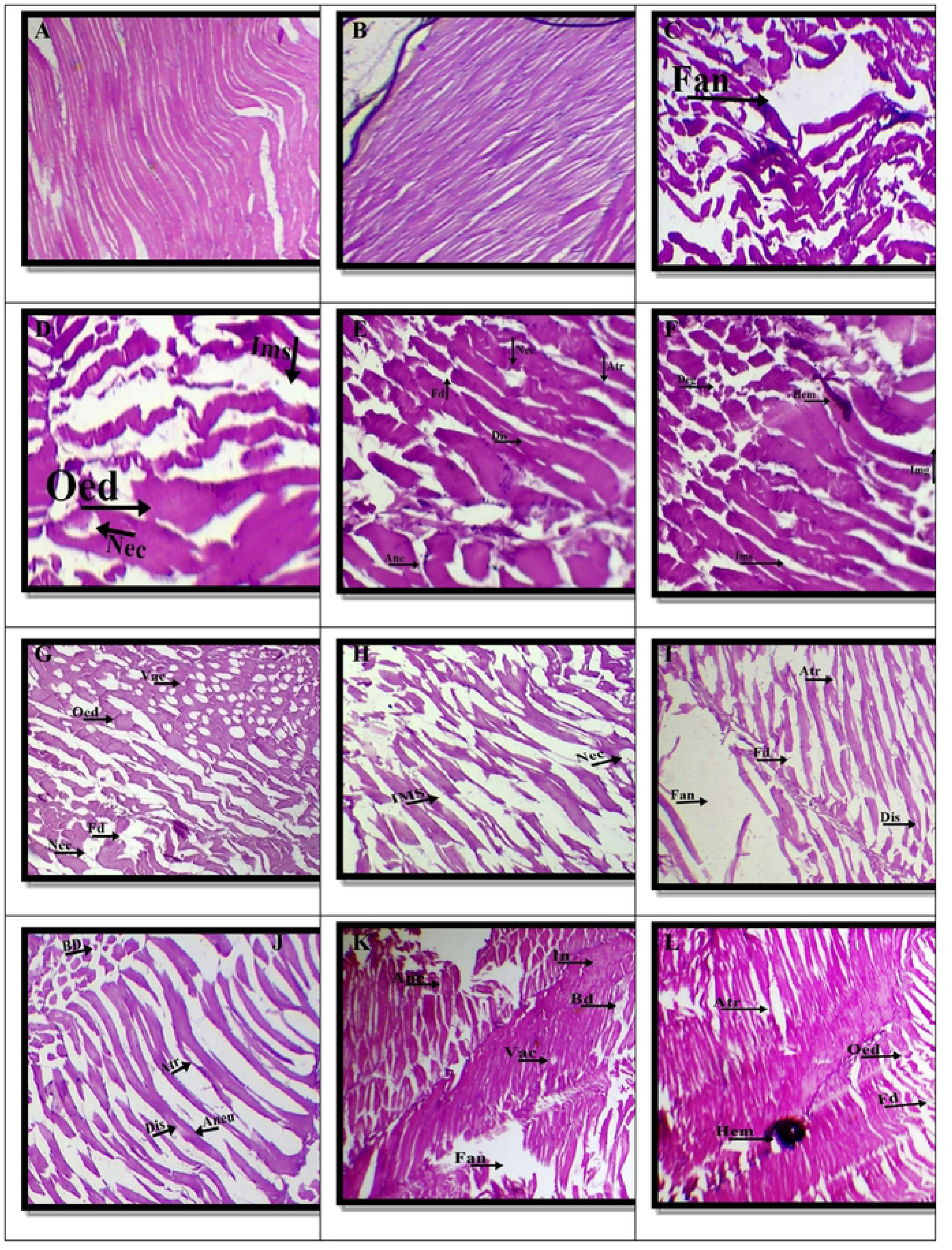
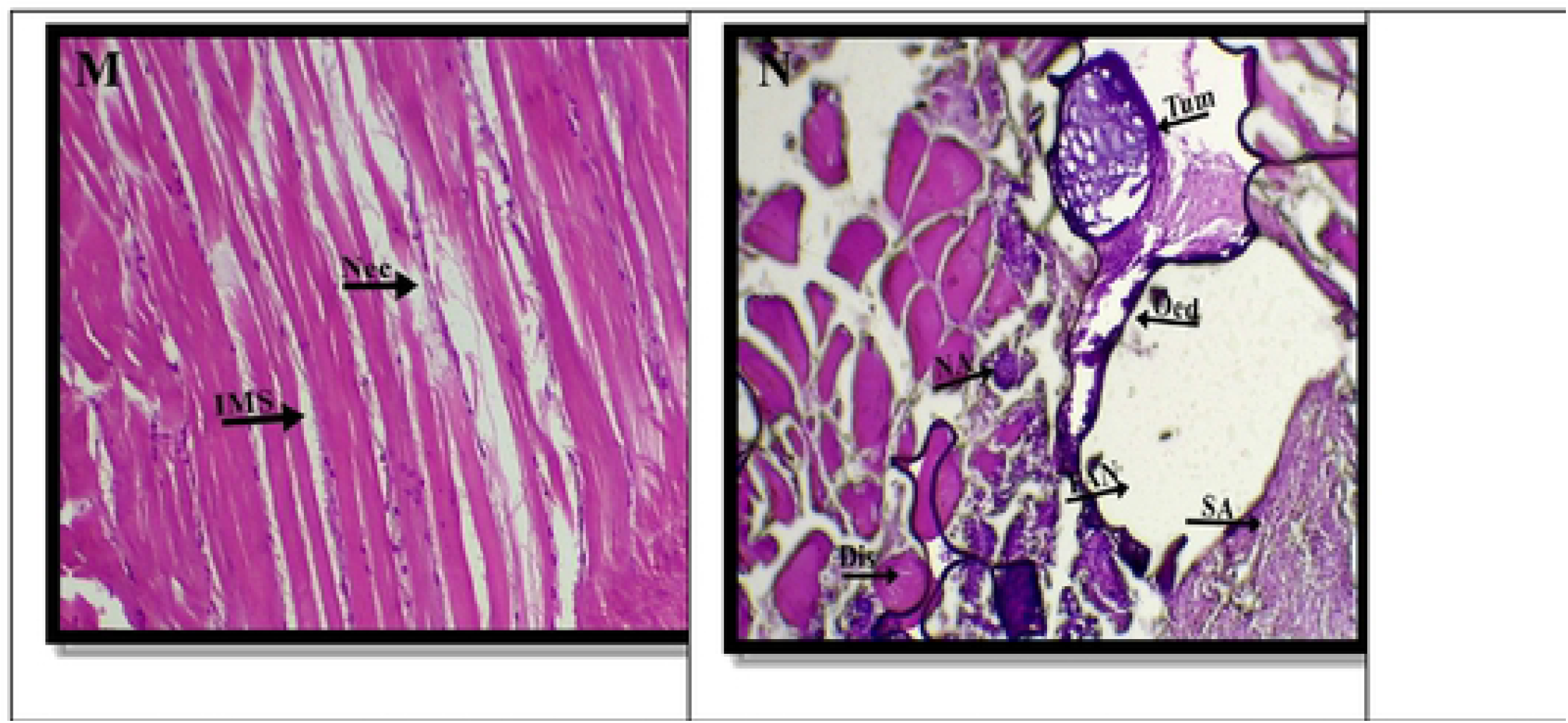
Histopathological alteration of transversely paraffin sectioned of muscle of *Oreochromis niloticus* electron micrographs (10×&40×) (H&E). (A&B) Control, (C-F) 1/3^rd^ of LC_50_, (G-J) 1/5^th^ of LC_50_, (K-N) 1/10^th^ of LC_50_. Normal architecture of muscle tissues were observed in the control fishes. **Abbriviations are as=** Hem=Hemorrhage, Oed=Oedema, Fd=Fiber damage, Aneu=Aneurysm, Atr=Atrophy, Bd=Bundle damage, Vac=Vacuolar degeneration, FAN=focal area of necrosis, Nec=Necrosis, IMS=Inter myofibril spaces, Dis=Disintegration, SA=Structural Alteration, NA=Nuclear Alteration, In=Inflammation and Tum=Tumor

Under both short and long term conditions, the heart of treated fish showed substantial alterations when compared to control fish, indicating that cardiac tissue is not resistant to this exposure (Table#2). The heart of an untreated O. niloticus was comprised of a mass of properly arranged cardiomayocytes. Furthermore, no evidence of necrosis could be found (Figure 3-A&B).

**Table 2:** Frequency and prevalence percentage (%) of histological alterations in the heart of *Oreochromis niloticus* exposed to sub-lethal levels of industrial effluents of Sunder Industrial Estate, Lahore.

| Histological Alteration | Control |  | Treatment Groups |  |  |  |  |  |
| --- | --- | --- | --- | --- | --- | --- | --- | --- |
|  | Group A |  | Group B |  | Group C |  | Group D |  |
|  | Frequency | % | Frequency | % | Frequency | % | Frequency | % |
| Hemorrhage | 0 | 0 | 1 | 3 | 3 | 10 | 5 | 16 |
| Oedema | 0 | 0 | 8 | 26 | 13 | 43 | 19 | 63 |
| Pericardium bending | 0 | 0 | - | - | 5 | 16 | 8 | 26 |
| Pericardium lifting | 0 | 0 | - | - | 4 | 13 | 5 | 16 |
| Aneurysm | 0 | 0 | 7 | 23 | 12 | 40 | 16 | 53 |
| Atrophy | 0 | 0 | 7 | 23 | 15 | 50 | 20 | 66 |
| Pericardium Damage | 0 | 0 | - | - | 2 | 6 | 5 | 16 |
| Vacuolar degeneration | 0 | 0 | 5 | 16 | 6 | 20 | 9 | 30 |
| focal area of necrosis | 0 | 0 | 4 | 13 | 10 | 33 | 16 | 53 |
| Necrosis | 0 | 0 | 5 | 16 | 18 | 60 | 13 | 43 |
| Hyperplasia | 0 | 0 | 2 | 6 | 8 | 26 | 16 | 53 |
| Leucocytes Infiltration | 0 | 0 | 1 | 3 | 2 | 6 | 8 | 26 |
| Myocarditis | 0 | 0 | 1 | 3 | 6 | 20 | 2 | 6 |
| Mean Value |  |  | 41 |  | 104 |  | 142 |  |

Affected myocytes in the treated groups showed symptoms of degeneration, including condensation, focal area of necrosis, vacuolar degeneration and loss of muscle striation, (Figure 3-G). Pyknosis or karyolysis was also occurring in a number of cells. Necrotic cells were found in the presence of inflammatory infiltrates (Figure 3-F&I). Ventricular lesions were the most prevalent, with mononuclear cells infiltrating both the spongy and compact layers (Figure 3-E).

Hemorrhage, atrophy, pericardium injury, myocarditis, necrosis, and other histopathological changes were observed in treated fish.. Maurya et al., 2019 (6) found significant alterations in heart tissue after exposure to industrial effluents, which is consistent with current study. An extensive myocarditis was a constant observation in all treatment groups; especially in group B (Figure 3-J) as compared to control group A, indicating inflammation. Inflammatory cells were seen in and around myocytes in a diffused or focused manner, with the compact layer showing the most inflammatory cells. In group B and C (Figure 3-D&G) respectively, leukocyte infiltrates were limited to the interface between the spongy and compact layers.

Cellular infiltration was broad and substantial in chronic exposure group D, with a significant number of necrotic myocytes. All of the treated groups showed hyperplasia, or hypergenesis, which is a rise in the quantity of organic tissue caused by cell proliferation. In chronic group D, there was a lot of hyperplasia (Figure 3-K). All treated groups had mild to moderate aneurysms, with stage I and II, in groups B, C, and D, respectively. In the myocardium of the treated groups, focal area of necrosis and hyperplasis were at stages II and III on the histological alteration index (Table 3). In addition, histological changes in the pericardium of the heart were documented i.e. pericardium bending, lifting and damaging as compared to control.

**Table 3:** Histological alterations index (HAI) calculated for determination of lesions in the Heart of *Oreochromis niloticus* exposed to sub-lethal level of industrial effluents of Sunder Industrial Estate, Lahore.

| Histologic Alteration | Importance factor | Index | Score Value (a) |  |  | HAI Value ( a×w) |  |  |
| --- | --- | --- | --- | --- | --- | --- | --- | --- |
|  |  |  | 1/3<sup>rd</sup><br>LC<sub>50</sub> | 1/5<sup>th</sup><br>LC<sub>50</sub> | 1/10<sup>th</sup><br>LC<sub>50</sub> | 1/3<sup>rd</sup><br>LC<sub>50</sub> | 1/5<sup>th</sup><br>LC<sub>50</sub> | 1/10<sup>th</sup><br>LC<sub>50</sub> |
| Hemorrhage | W<sub>HC1</sub>=1 | I<sub>HC</sub> | 1 | 1 | 1 | 1 | 1 | 1 |
| Oedema | W<sub>HC2</sub>=1 |  | 2 | 2 | 2 | 2 | 2 | 2 |
| Pericardium bending | W<sub>HR1</sub>=1 | I<sub>HR</sub> | - | 1 | 2 | - | 1 | 2 |
| Pericardium lifting | W<sub>HR2</sub>=1 |  | - | 1 | 1 | - | 1 | 1 |
| Aneurysm | W<sub>HR3</sub>=1 |  | 1 | 2 | 2 | 1 | 2 | 2 |
| Atrophy | W<sub>HR4</sub>=2 |  | 1 | 2 | 2 | 2 | 4 | 4 |
| Pericardium Damage | W<sub>HR5</sub>=1 |  | - | 1 | 1 | - | 1 | 1 |
| Vacuolar degeneration | W<sub>HR6</sub>=2 |  | 1 | 1 | 2 | 2 | 2 | 4 |
| focal area of necrosis | W<sub>HR7</sub>=3 |  | 1 | 2 | 2 | 3 | 6 | 6 |
| Necrosis | W<sub>HR8</sub>=3 |  | 1 | 2 | 2 | 3 | 6 | 6 |
| Hyperplasia | W<sub>HP1</sub>=2 | I<sub>HP</sub> | 1 | 2 | 2 | 2 | 4 | 4 |
| Leucocytes Infiltration | W<sub>HP1</sub>=2 | I<sub>HI</sub> | 1 | 1 | 2 | 2 | 2 | 4 |
| Myocarditis | W<sub>HP2</sub>=1 |  | 1 | 1 | 1 | 1 | 1 | 1 |
| Mean value of HAI<sub>Kidney</sub> |  |  |  |  |  | 19 | 33 | 38 |

Essien and Chinwe (33), observed similar results after exposing *Oreochromis niloticus* to effluents. After 96 hours of sublethal exposure to malathion, Mager and Dube (34) found structural changes in the heart muscle of *Channa puncctatus*, including atrophy and congestion. After toxicant exposure, Das and Mukherjee (35) found significant pericardial thickening and leucocyte infiltration in the *labeo rohita*.

In all treated groups, however, kidney tissues of O. niloticus revealed expanded bowmen space and dilated glomerulus (Figure 4-C, H, and N). Group B kidney tissues displayed necrotic proximal tubules, damaged renal/glomerular tubules, and hemorrhage after exposure to 1/3rd of LC50. The kidney of O. niloticus in the control group, on the other hand, had a normal histological features, with roughly spherical glomeruli and appropriate bowman space (Figure.4-A). Normal structure was seen in the lumen of distal tubules and the brush border of proximal tubules and (Figure 4-B).

As demonstrated in (Table 4), frequency and prevalence percentage of kidney lesions increased with an increase in exposure length to 6 months compared to 3 and 15 days. Damage to the nephronic and tubular levels of the kidney has been discovered in a recent study (Figure 4). Hemorrhage and vacuolization (stage I damage) in group B, renal and tubular degeneration, (stage II damage) in group C, and proximal tubular necrosis (stage III damage) in group D were all observed. Corpuscles showed obvious damage, including glomerulus shrinkage, glomerular structural alteration had same degree of damage, and increase in glomerulus space (stage I, II and III damage) showed duration dependent alteration in all treated groups (Table 5). Other histological abnormalities associated with blood cell infiltration were also seen in the current investigation. In fish, renal tubule necrosis impacts metabolic activity and promotes metabolic disorders (36).

**Table 4:**
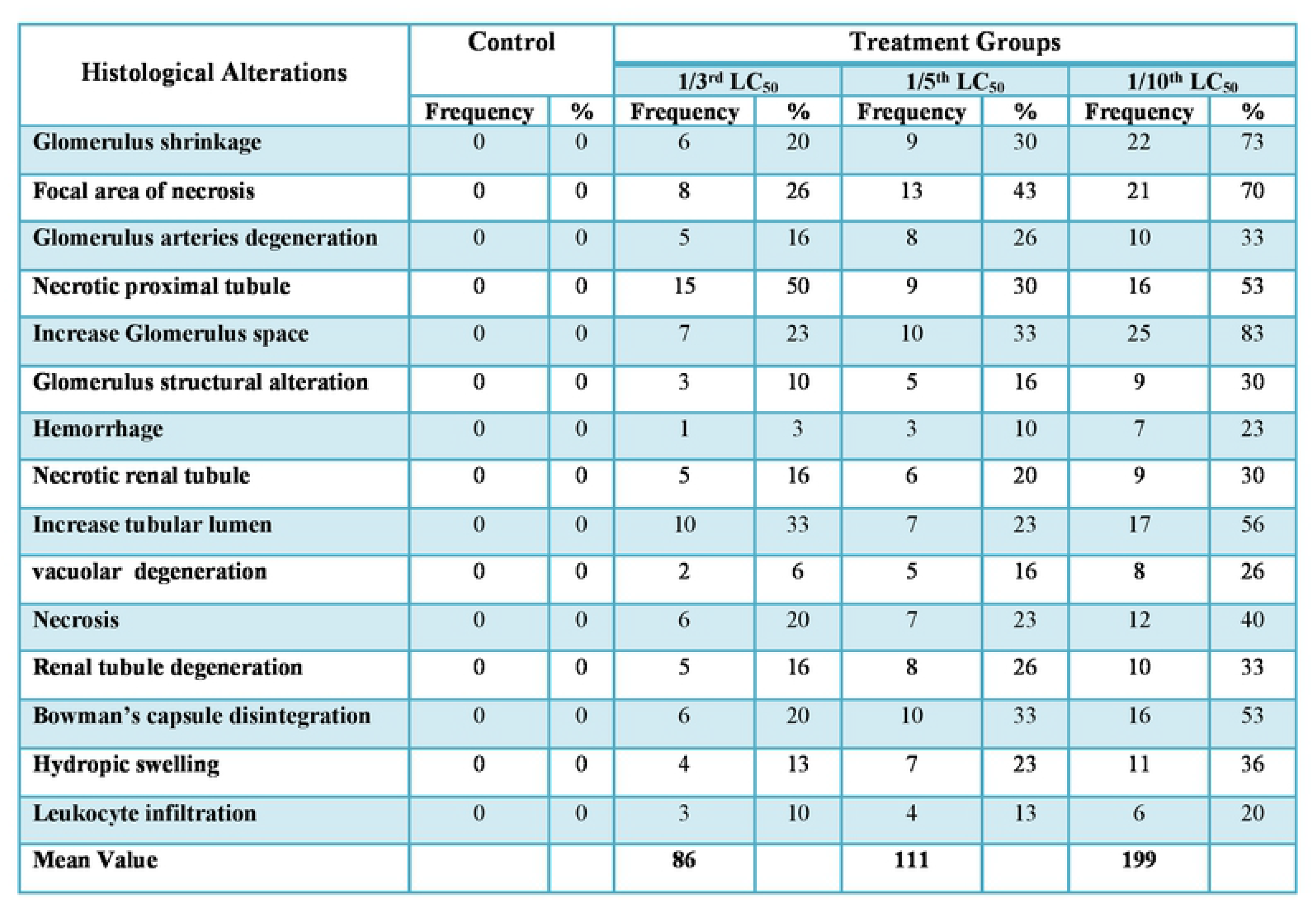
Frequency and prevalence percentage (%) of histological alterations in the Kidney of *Oreochromis niloticus* exposed to sub-lethal level of industrial effluents of Sunder Industrial Estate, Lahore.

**Table 5:**
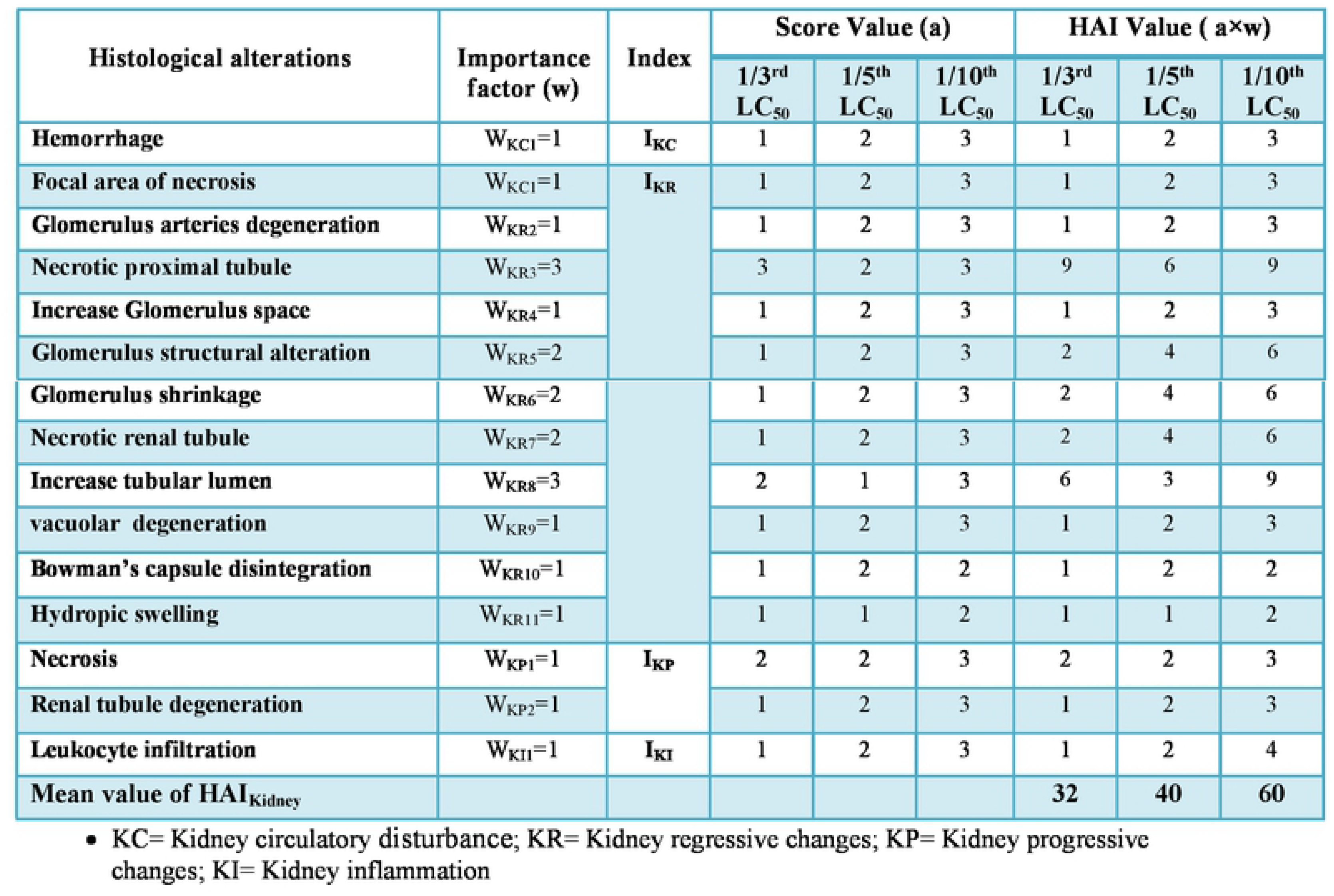
Histological alterations index (HAI) calculated for determination of lesions in the Kidney of *Oreochromis niloticus* exposed to sub-lethal level of industrial effluents of Sunder Industrial Estate, Lahore.

The degenerative process contributes to tissue necrosis with chronic exposure. So in group D, the maximum prevalence of histopathological alteration of all three categories (stages I, II, and III) was seen. When compared to changes observed after 3 and 15 days, the damage was much more severe. The tubular epithelium showed necrosis in renal tubules with focal areas of necrosis (Figure 4-N) with deshaped glomerulus (structural alteration) (Figure 4-L&M). Degeneration, vacuolization, and disordered capillaries in the glomerulus were detected in all treatment groups (Figure 4-D, E, G, and N).

In the exposed fish, structural damage varied from mild (3 days), moderate (15 days) and severe (6 months). The identified degree of tissue alteration values is consistent with previous studies that treated *Labeo rohita* with chronic wastewater exposure, as reported by (11). Shrinkage, destruction of tubular epithelium, vacuolization, and glomerulus injury are some of the other histological abnormalities seen in the kidney. Navaraj and Yasmin (37) found similar results in *Oreochromis mossambicus* subjected to tannery industry waste water and *Rasbora daniconius* exposed to sub-lethal concentrations of industrial effluents (38).

Histopathology of transversely sectioned muscle tissues of *O. niloticus* exposed to varying concentrations of LC_50_ revealed different histological alterations in duration dependent manner, including oedema (Figure 5-D&G), muscle fiber necrosis (Figure 5-H) and inflammation (Figure 5-K), enlarging lesions in muscular tissue’s epidermis (Figure 5 J&K), and many others (Table 6). Muscle fibers with a wispy coating of areolar endomysial connective tissue that encased muscle fibers were seen in control tilapia tissues (Figure 5-A). Many thick and thin alternating myofilaments were observed in myofibrils with a diameter of 1 micrometer (Figure 5-B).

**Table 6:** Frequency and prevalence percentage (%) of histological alterations in the Muscle of *Oreochromis niloticus* exposed to sub-lethal levels of industrial effluents of Sunder Industrial Estate, Lahore.

| Histological Alteration | Control |  | Treatment Groups |  |  |  |  |  |
| --- | --- | --- | --- | --- | --- | --- | --- | --- |
|  | Group A |  | Group B |  | Group C |  | Group D |  |
|  | Frequency | % | Frequency | % | Frequency | % | Frequency | % |
| Atrophy | 0 | 0 | 9 | 30 | 15 | 43 | 20 | 66 |
| Oedema | 0 | 0 | 7 | 23 | 14 | 46 | 18 | 60 |
| Hemorrhage | 0 | 0 | - | - | 1 | 3 | 7 | 23 |
| Fiber damage | 0 | 0 | 9 | 30 | 12 | 40 | 17 | 56 |
| Aneurysm | 0 | 0 | 8 | 26 | 11 | 36 | 16 | 53 |
| Inflammation | 0 | 0 | 5 | 16 | 8 | 26 | 11 | 36 |
| Bundle damage | 0 | 0 | 6 | 20 | 10 | 33 | 15 | 50 |
| Vacuolar degeneration | 0 | 0 | 4 | 13 | 6 | 20 | 10 | 33 |
| focal area of necrosis | 0 | 0 | 5 | 15 | 12 | 40 | 18 | 60 |
| Necrosis | 0 | 0 | 6 | 20 | 22 | 73 | 16 | 53 |
| Inter myofibril spaces | 2 | 6 | 10 | 33 | 18 | 60 | 25 | 83 |
| Disintegration | 0 | 0 | 8 | 26 | 15 | 50 | 20 | 66 |
| Tumor | 0 | 0 | - | - | - | - | 3 | 10 |
| Structural Alteration | 0 | 0 | - | - | - | - | 10 | 33 |
| Nuclear Alteration | 0 | 0 | - | - | - | - | 7 | 23 |
| Mean Value | 2 |  | 77 |  | 144 |  | 215 |  |

The tissue sections of fish subjected to a chronic dosage of effluents revealed significant alterations. Disintegration (Figure 5-N) in muscle bundles along with focal areas of necrosis were among the pathological findings (Figure 5-K). In addition, group D had moderate vacuolar degeneration in muscle bundles, but group B had significant vacuolar degeneration and muscle bundle atrophy (Figure 5-L). There was oedema between muscle bundles in group D, as well as muscle fiber breaking. The histopathology of the muscle of the treated fish revealed gradual damage to the structure of the muscle when the effluents were applied for longer periods of time. As the effluent duration increased, Nagarajan and Suresh (39) reported comparable outcomes in the muscle tissue of the fish *Cirrhinus mrigala*.

The muscle layer that made up skeletal muscle in normal fish was the lateral muscle layer. Whereas cell fading was observed in the infected muscle of groups B and C (Figure 5-D&G), necrosis was observed in the infected muscle of group A. In addition, the damaged muscle generated by the 1/3rd and 1/5th doses of industrial effluents in groups B and C, which was characterized by muscular abnormalities, can be seen in (Figure 5-E&I). Inflammatory cell infiltration is a pathogenic reaction that occurred in all of the treatment groups.

Thickening and intermyo fibril spaces (Figure 5-M), necrosis (Figure 5-L), hemorrhage, and lesions with reduced compactness were detected after chronic exposure of effluents in group D. While sub-chronic concentrations of effluents caused pronounced intramuscular oedema and mild dystrophic changes in group C. Damaged myofibrils (Figure 5-G) and bundles ((figure 5-I) and formation of gap between muscle bundles were noted as significant changes, which eventually led to degeneration of muscular bundles along focal areas of necrosis, and atrophy (Figure 5-J).

Muscle tissue in the chronic exposure group D showed dystrophic alterations. The myoseptum, which separates each myotome, appears to have vanished from the muscle. In the muscle portion, there is also disintegrated epidermis. Intramuscular oedema is a common symptom, with the highest incidence in group D and the lowest in the other treated fish groups. Furthermore, chronic exposure resulted in the formation of tumor cells (Figure 5-M), as well as other serious changes. Histological alteration index (HAI) of muscle for group D showed highest alteration as compared to group B and C (Table 7).

**Table 7:**
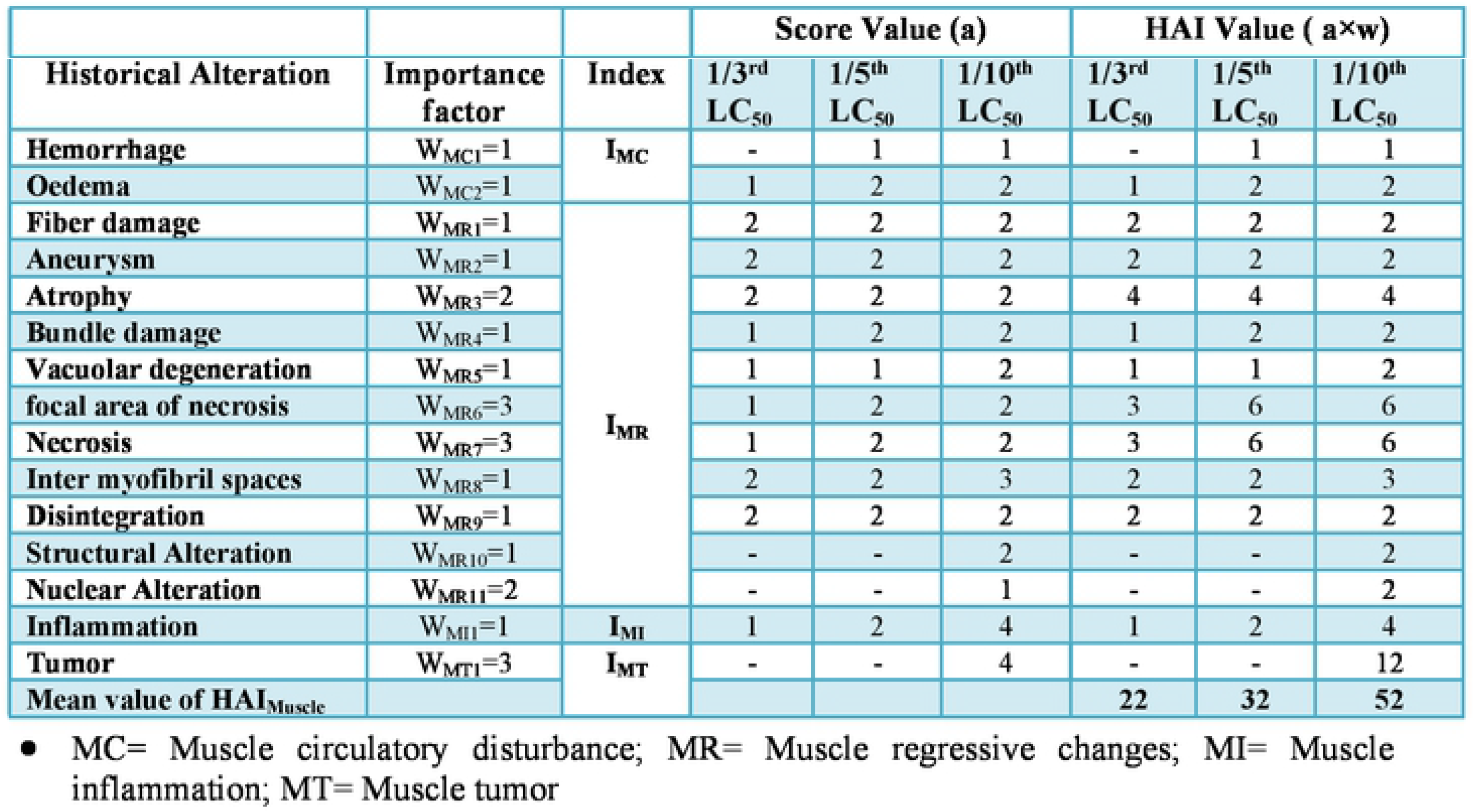
Histological alterations index (HAI) calculated for determination of lesions in the Muscle of *Oreochromis niloticus* exposed to sub-lethal level of industrial effluents of Sunder Industrial Estate, Lahore.

Minor lesion, inclusion of bodies and necrosis, cellular degeneration and inflammation, were detected in the muscular tissue of the fish *O. niloticus* by Oluwatoyin (40). Similarly, Nagarajan (30) found that when effluent concentrations increased, muscle tissue began to deteriorate. Many researchers have investigated the effect of various contaminants on the muscles of fish have identified histological abnormalities in the muscles of the analyzed fish (41); (42) and (36).

Pollution is one of the most damaging human influences on the aquatic environment, resulting in chronic stress conditions that harm aquatic species. Many researchers have studied the effects of various pollutants on various fish organs, and the current findings are consistent with their findings (33) and (43). The presence of histological alteration in several organs of fish has been analyzed, revealing that these effluents cause serious damage to muscle, kidney, and heart. The histological studies are useful tool for determining the effects of numerous anthropogenic poisons on living organisms. In a biological system, histopathological biomarkers reflect the population’s overall health (44).

The current investigation found that the water quality in Sunder Industrial Estate had deteriorated, with multiple physicochemical characteristics exceeding the standard discharge limits. The lesions found in histological examinations revealed cellular and organ damage, as well as confirming *Oreochromis niloticus* sensitivity to industrial effluent. The fish’s histological modifications show how they deal with the stress of inadequate water quality. Present study revealed that *Oreochromis niloticus* should be deployed as sentinels in water quality monitoring programs.

## Conclusion

Histopathological data revealed that a composite of industrial effluents was capable of altering the gross structure of fish organs, resulting in increasing degrees of tissue damage in correlation to the effluent dose duration. In addition, important physico-chemical parameters of industrial waste water were also recorded which were beyond the stander range.

## Acknowledgment

I am heartily thankful to my supervisor for her fruitful guidelines for this research and Higher Education Commission Pakistan too. This research was funded by Higher Education Commission Pakistan under the project#7423/KPK/NRPU.

